# In-silico studies on structural and thermodynamics basis of interaction of lac repressor protein binding to different DNA operators

**DOI:** 10.1101/2024.07.07.602393

**Authors:** Soumi Das, Siddhartha Roy, Dhananjay Bhattacharyya

## Abstract

Transcription factors, which bind to a large number of sequences in the sea of genome non-specifically, affect the final outcome in a sequence-dependent manner and intrigued us to explore whether there exists any conformational preference of the DNA. Lac-repressor, binds to major groove of DNA with strong sequence specificity, also interacts with the minor groove of its operator by partially intercalating two symmetry related Leucine side-chains between two CG base pairs. We attempt to elucidate mechanisms of the structural and dynamic alterations of two different Lac operator DNA sequences upon Lac repressor recognition using molecular modelling and all-atom molecular dynamics simulations. The difference of hydrogen bonding network and deformation of DNA base pair step produce the asymmetric dynamics of the two subunits of Lac repressor protein. Variation of minor groove widths can direct overall shape complementarily between the DNA and protein surfaces indicating DNA is significantly deformed to accommodate the protein fold. Bending flexibility indirectly modulates the Lac binding activity that controls transcription efficiency. Thus we suggest that Lac operator is unique as it uses both Lock & Key mechanism for DNA recognition, where protein induced structural alteration of the receptor DNA is not required, and also Induced Fit/Conformational Selection model of recognition.

**Graphical Abstract:** 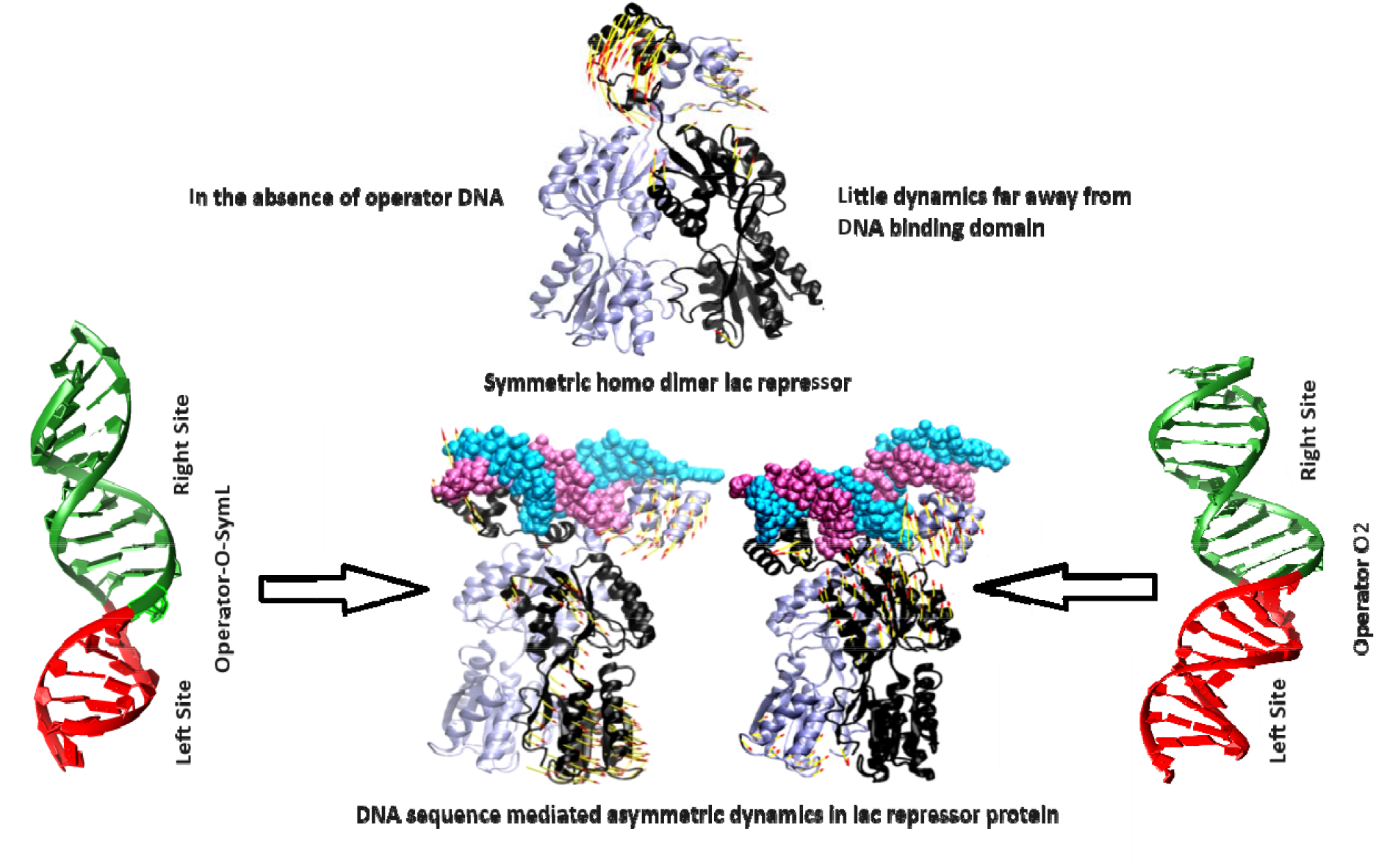

**Highlights:**

- Recognition of two different DNA Sequences by prokaryotic transcription factor lac repressor is modulated through asymmetric protein dynamics that have emerged in terms of conformational entropy and essential dynamics.
- The difference of the hydrogen bonding network and the deformation of the DNA base pair step produces the asymmetric dynamics of the two subunits of the lac repressor protein.
- The difference in overall protein dynamics upon interaction with DNAs of different sequences is the key feature in stabilizing the gene regulatory networks differently.

## 1. Introduction

The mechanism by which genetic regulatory proteins discern specific target DNA sequences remains a fascinating world of biophysics **[1, 2]**. There are numerous systems in different forms of life, prokaryote as well as eukaryote, where such DNA sequence-specific binding to different gene regulatory proteins takes place. Significant experimental data is available for some of them, such as gal, lac, cro, etc.

The Lac repressor has two different effectors: operator DNA and inducer. Thus, Lac repressor is a heterotropic system. Lac repressor, a tetramer of four identical subunits, normally binds tightly to a specific region in the bacterial DNA, termed as the operator, which is next to a region that encodes three lactose-metabolizing proteins and inhibits the production of the Lac proteins by occluding the RNA polymerase binding or by prompting DNA looping. But in the presence of lactose, the Lac repressor binds to allolactose and induces an allosteric change in the Lac repressor protein. As a result, the Lac repressor is no longer able to bind to the DNA (cognate operator). Then, RNA polymerase is free to transcribe the gene, and the lactose-metabolizing proteins are made. Catabolite Activator Protein (CAP), known as c-AMP response protein (CRP), binds to c-AMP and promotes the transcription of lactose-catabolizing genes. Two distinct DNA binding factors (a) c-AMP-CRP complex acts as a positive regulator and (b) Lac repressor acts as a negative regulator of the Lac operon. Transcription factors form a notable class of proteins where allostery plays an important role. A few cases have been reported in which the DNA sequence itself plays the role of an allosteric ligand and changes the functional outcome in terms of gene regulation. However, little is known about the role of dynamics in such situations. Alterations in overall protein dynamics upon interaction with the different operators have crucial importance in the gene regulatory network. Transcription factors, which bind to a large number of sequences in the sea of the genome in a non-specific way and affect the outcome in a sequence-dependent manner, intrigue us to explore how two distinct DNA sequences affect protein dynamics **[3, 4]**.

We use the *Escherichia coli* lactose repressor (LacI) as a model protein and its two operator target sequences O2 and SymL as effectors shown in **Figure 1 (a)**. The Lac repressor comprises of four 4 indistinguishable monomers of 37.5 kDa each. Structurally it can be viewed as a dimer of dimers. Each monomer consists of (a) the amino-terminal headpiece (HP) (residue numbers 1-62) which is capable of DNA operator recognition and binding, (b) the core domain (residue numbers 63-329) which encompasses the inducer binding site and dimerization interface, and (c) the carboxy-terminal tail (residue numbers 330-360) which is responsible for tetramer formation. The core consists of two subdomains: the N-subdomain, containing residues 63-161 and 293-320, and the C-subdomain, with residues 162-289 and 321-329 (**Figure 1(b)**). The hinge helix communicates an allosteric signal from the DNA Binding Domain to the Core Domain. This crystal structure shows the four monomers (colours red, green, yellow and cyan) bound to two DNA operators (gray).

**Figure 1(a).**
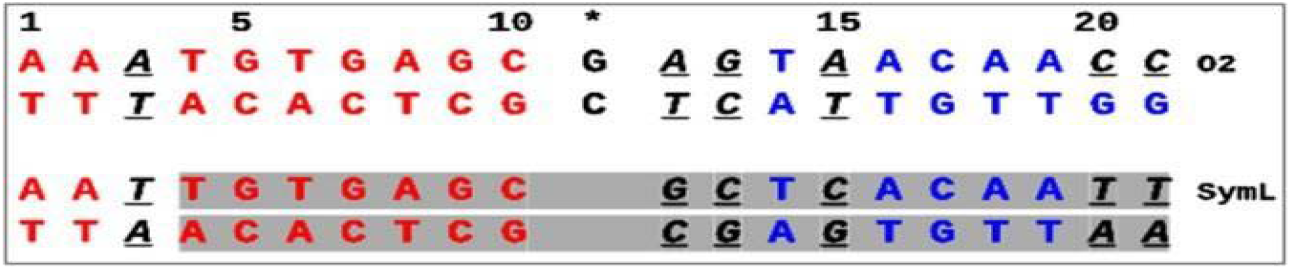
Sequences of naturally occurring pseudo-palindromic O2 and palindromic, symmetric, and synthetic SymL lac operators. The asterisk denotes the central base pair. The bases conserved in operators are highlighted in red and blue at the left and right sites respectively. The two binding sites within the natural operator are referred to as the left (basepairs 1-10) consensus and right non-consensus (12-21). In operator O2, the asymmetric regions between the two sites relative to the central base pair are underlined, italic, and black colored with respect to the SymL operator. The SymL operator (ideal) is a palindrome of the left half-site of theO1 sequence with the central GC base pair deleted.

**Figure 1(b).**
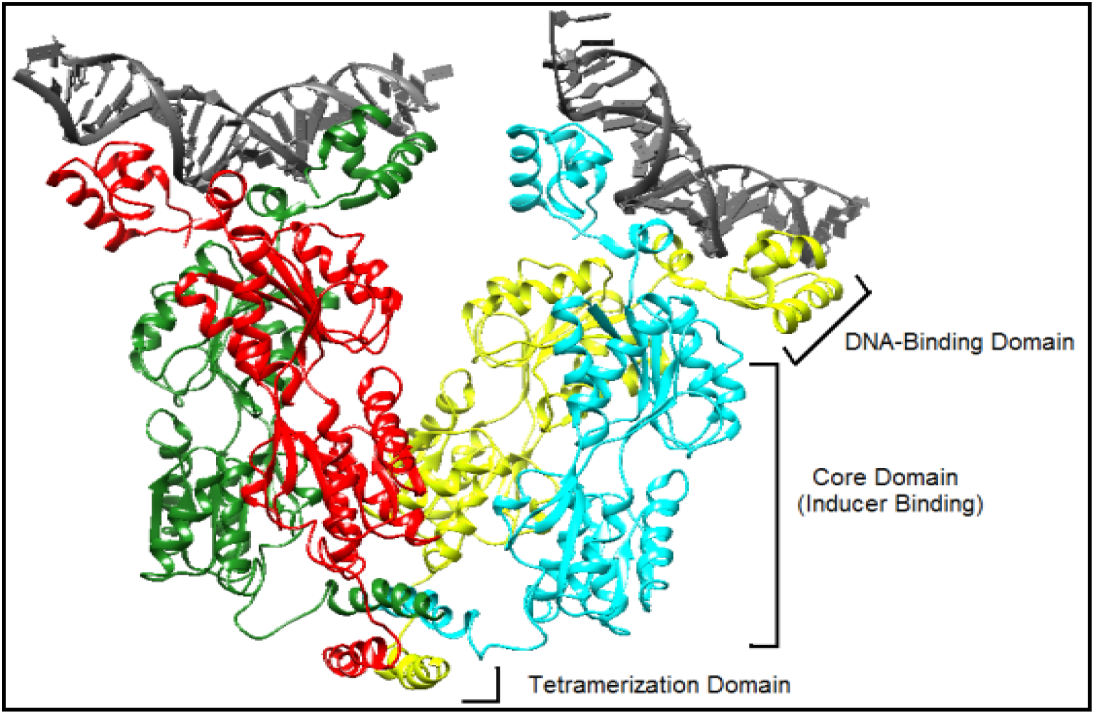
Ribbon Representation of a 4.80Å Crystal Structure of Tetrameric-Lac repressor Bound to Two DNA Operators (PDB-ID-1LBG) using Chimera.

We attempt to address whether is it true that modulation of protein dynamics by the different DNA homo-dimer lac repressor is symmetric. We shed light on how two different operator DNA sequences O-SymL and O2 alter protein dynamics of lac repressor protein differently using molecular dynamics simulation of repressor-operator complexes, free operator O-SymL, free operator O2, free lac repressor protein systems. A detailed microscopic insight into recognition mechanism of lac repressor-operator complex would be helpful to understand conformational heterogeneity of protein-DNA complex elaborately.

## 2. Method

### 2.1 Model Building

The coordinates of the Lac dimer and Lac bound DNA were obtained from the Protein Data Bank **[5]** having PDB ID 1EFA which contains the whole Lac repressor protein as dimer. Chains A and B of protein were considered. Chain C containing residues 46-331 was removed to reduce the molecular size making the repressor fully symmetric. Anti-inducer ONPF binding to repressor-operator complex has no significant influence on the structure of ligand binding pocket of Lac as well as the conformation of the Lac-dimer and DNA. Therefore, we have deleted ONPF and crystal waters. This is the model of Lac-O-SymL repressor-operator complex.

Unfortunately, there is no X-ray crystallographic or NMR structure of Lac-O2 repressor-operator complex containing the intact Lac repressor protein consisting of two subunits (residues 2-329) of DNA binding and core domains. In order to model Lac-O2 complex, we superpose the N-terminal domain of the protein dimer of 1EFA on the N-terminal DNA binding domain of the protein of the NMR derived structure 2KEJ. Further, the existing DNA binding domain of 2KEJ was removed. This derived protein-DNA complex is considered as a model of protein-DNA complex for Lac-O2 system. The 8^th^ model from NMR ensembles of 2KEJ was identified as closest to the average NMR structure by WHAT IF server. The protein–DNA contacts are retained by superposing N-terminal DNA binding region of the Lac repressor protein in the two structures. The modelled Lac-O-SymL and Lac-O2 complex were further subjected to energy minimization by steepest descent with 5000 steps and conjugate gradient with 10,000 steps, using CHIMERA **[6]** in order to remove steric clashes generated during structure modelling. The free DNA models were obtained from 1EFA and 2KEJ by removing the protein molecules. In the Lac-O2 complex, chain A protein subunit binds at consensus half and that of chain B interacts with non-consensus half.

### 2.2 Molecular dynamics simulations

MD simulations with explicit solvent for both protein-DNA complexes (Lac-SymL and Lac-O2 complex), protein unbound free operator (SymL and O2) DNA and free protein (operator unbound) **(Table 1)** were performed with GROMACS 5.1 **[7]**, using an improved Amber-ff99SB-ILDN **[8]** force field corresponding to neutral pH for 300 ns each. In case of His residues, the Nε is considered as secondary amino nitrogen and Nδ as imino nitrogen. In order to confirm the effect of force field on the structural properties obtained from MD simulation, we have performed an additional simulation of the SymL-protein-DNA complex with a more recent force field named Amber-ParmBSC1 **[9]**. The systems were solvated with TIP3P water model in cubic box using a distance of 1 nm between the complex and the edge of the box. The LINCS algorithm was utilized to constrain all bonds involving hydrogen atoms. The Particle Mesh Ewald (PME) method **[10]** was used to evaluate long range electrostatic interactions. Short-range distance cut off for non-bonded interactions was set to 0.9nm. The Protein-DNA complex systems ((Lac-O-SymL) and (Lac-O2) approximately contain 165000 and 179000 atoms respectively and Protein unbound operator DNA ((O-SymL) and (O2)) approximately contains 50000 and 80000 atoms respectively) were equilibrated under constant temperature (300 K) and pressure (1 bar) for 300 ps, using the Parrinello-Rahman method **[11]**. The production runs are performed for 300 ns each under NPT ensemble with 2 fs time step. The atomic coordinates were saved every 10.0 ps, and thus, 30,000 structures were collected with each system for further analysis.

**Table 1:**
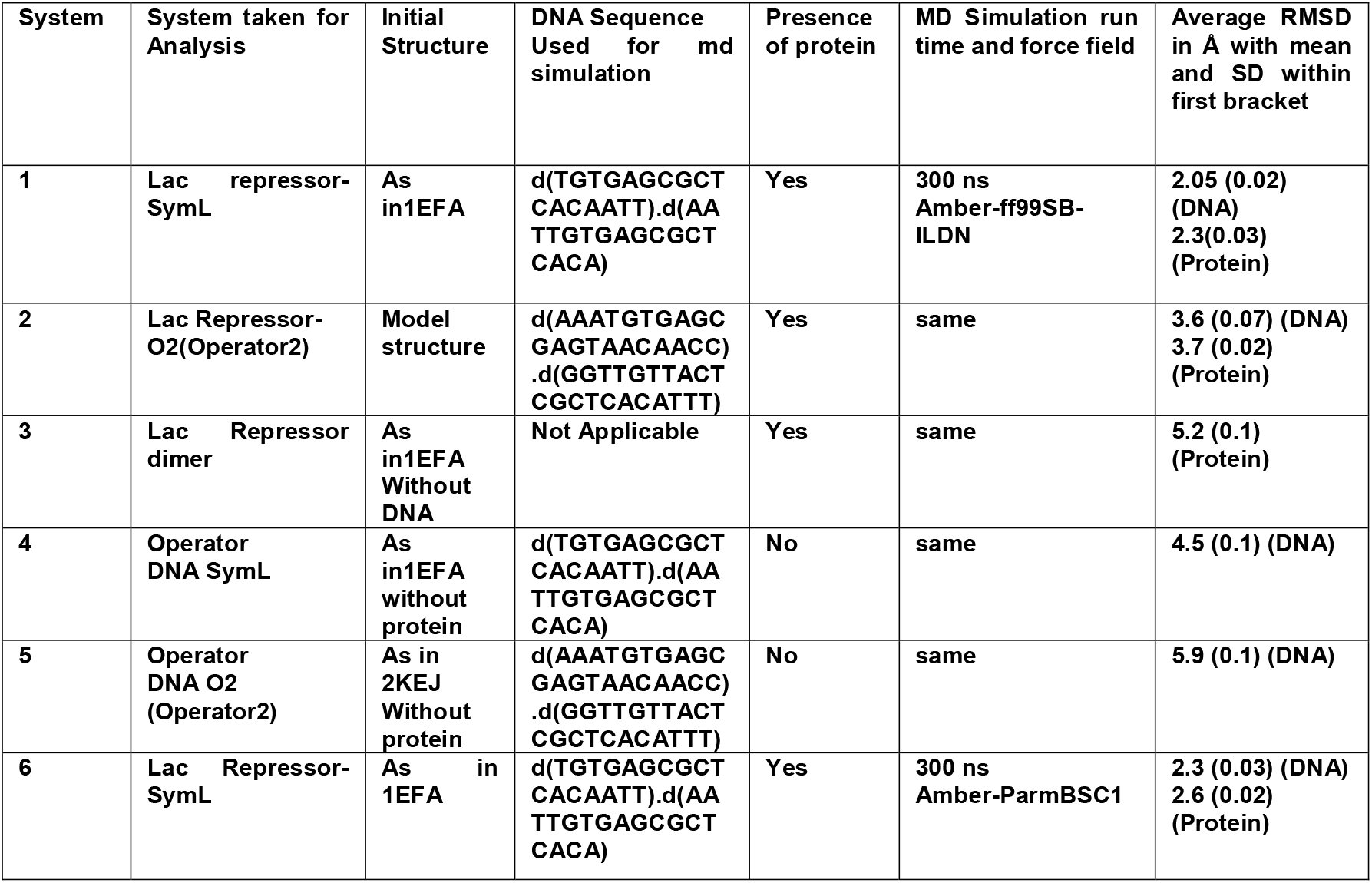
System taken for study using MD.

### 2.3 Structure analysis

Two DNA strands and two protein subunits were apparently found to be separated quite often due to periodic boundary condition (PBC) imposed during simulation. The same incident happens for every system, which could not be corrected by “PBC MOL” option of GROMACS or wrap option of VMD. We resolved the problem by converting MD trajectory of each system into PDB format using GROMACS utility “trjconv” and running in-house FORTRAN program on the corresponding PDBs to wrap up the separated strands for correct analysis. Root mean square displacement (RMSD) and root mean square fluctuation (RMSF) analyses of the trajectories were performed using GROMACS after the above operations. Detection of amino acid residues in alpha-helical, beta-sheet or turn conformations were done using GROMACS that follows DSSP **[12]** algorithm for assignment of secondary structure. The hydrogen bonding information between protein and DNA during MD runs were obtained by a modified version of pyrHBfind **[13]** using H-bond distance cut off 3.0 Å and angle greater than 150°. Side chain torsion angles, chi 1(χ1) of the amino acid residues were studied using VMD.

The global bending of the DNA double helix upon complex formation for Lac-O-SymL and O2 operators were estimated using NUPARM in terms of bending angle for which two average helix axes were fitted to the DNA systems by considering C1’ atoms of the non-terminal base pairs on either side of the junction and excluding those present near the terminal due to fraying effect. The helix-axis determination algorithm follows that adopted by Dickerson in the FREEHELIX program **[14]. Figure 3**. clearly sheds light on the stretch of two helix axes for bending angle calculation. The details of the helix-axis chosen for the DNA systems are mentioned in **Table 2**.

**Table 2:** Stretch of Helix axis.

| System | Stretch of helix axis 1 | Stretch of helix axis 2 |
| --- | --- | --- |
| Operator O-Sym | Chain D: 6-11 (Strand 1) | Chain D: 14-20 (Strand 1) |
|  | Chain E: 17-12 (Strand 2) | Chain E: 9-3 (Strand 2) |
| Operator O2 | Chain C: 1-9 (Strand 1) | Chain C: 12-21 (Strand 1) |
|  | Chain D: 21-13 (Strand 2) | Chain D: 10-1 (Strand 2) |

### 2.4 Study of conformational thermodynamics (entropy) by histogram-based method (HBM)

Conformational entropy of the systems was calculated following the Gibbs entropy formula. The conformational entropy *s*_*conf*_(χ) for a particular side chain dihedral angle, χ1 is

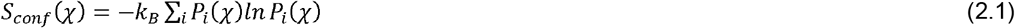

Where the sum is taken over the histograms of bins i with nonzero values of probability distribution function *P*_*i*_.

We extract the side chain conformational entropy from the histograms of the side chain dihedral angle χ1 **[15]**. We generate a histogram of χ1 for each residue within 0° to 360° and divide it into 90 equal bins. The change in side chain conformational entropy for χ1 due to binding of the repressor protein with operator DNA is

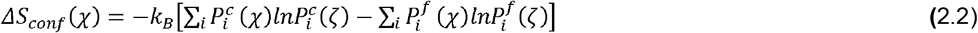

The conformational changes of DNA structure were studied using inter base pair parameters and inter base pair step parameters by NUPARM software **[16, 17]**.

### 2.5 Principal Component Analysis (PCA)

Reducing the dimensionality of the huge data obtained from molecular dynamics simulations can help in identifying configurational space that contains only a few degrees of freedom in which harmonic motion persists. Principal Components Analysis (PCA) method is based on the construction and diagonalization of the covariance matrix (C_ij_) of complex sets of variables and is used to reduce the higher-dimensional data to extract meaningful information from the protein-DNA complex throughout the simulation. The elements C_**ij**_ in the matrix are defined by **C**_**ij**_ **= < (x**_**i**_**-<x**_**i**_**>) (x**_**j**_**-<x**_**j**_**>) >** where <x_i_>, <x_j_> are the mass-weighted mean Cartesian coordinates of the atoms present in the system and x_i_ (or x_j_) is the coordinate of the i^th^ (or j^th^) atom of the systems, and “< >“ indicates ensemble average. The eigenvectors represent the direction of the coordinated motion of atoms, and the eigen values represent the magnitude of the motion along the movement direction. Usually, the first few principal components (PCs) describe the most important slow modes of the system, which are related to structural changes of the complex observed during the simulation. The GROMACS program gmx covar is used to calculate and diagonalize the mass-weighted covariance matrix. The generated eigenvectors are analysed using the program gmx anaeig in the GROMACS program suite. The phosphate atoms of DNA and Cα atoms of protein are only used to construct the covariance matrix, since the size of the matrix varies with the square of the number of atoms for which the covariance is calculated.

A collective ensemble of conformational fluctuations involving multiple regions of a biological macromolecule is more commonly referred to as breathing motions. 1^st^ and last frame mean the two end points of vibration as obtained from PCA are designated as PC1 1^st^ and PC1 last. In-house FORTRAN program, was used for Porcupine plot analysis requires two pdb files from MD trajectory indicating two end points to draw the vectors which refer to the direction and extent of the motion.

The Pymol, UCSF Chimera and Visual Molecular Dynamics (VMD) packages are used for visual assessment of the trajectory files and to generate images. Graphs are generated by gnuplot, MS-excel and MATLAB. MD simulations are performed using a Dell Power Edge server with a CentOS6 GNU/Linux operating system.

## 3. Result and Discussion

### 3.1 Evaluation of the convergence behaviour of the MD Simulations

The equilibrated snapshots at 300 ns time span of lac repressor unbound free operator DNA O-SymL and O2 along with lac repressor-O-SymL complex, lac repressor-O2 complex systems were shown in **Figure 3**. Structural stability and conformational variability were measured by root mean square deviations (RMSDs) of the Cα atoms of dimeric protein in operator bound and unbound state relative to the initial structure **(Figure S1)**. The RMSDs of the DNA-bound system are significantly lower than those of the free protein chain. It indicates that the Lac repressor-SymL and Lac-repressor-O2 (Operator2) are well constrained in the DNA-bound system. The RMSD of Cα atoms for all amino acid residues of both Lac-O2 and Lac repressor-SymL indicates that the systems reach the equilibrium state at 50 ns. A jump in RMSD is observed within the first 10 ns is a consequence of relaxation of the starting model. To explore the effect of target DNA binding on the protein dynamics, the free protein dimer is also subjected to molecular dynamics simulations. The RMSD values of the free protein (operator unbound) and free DNA systems continue to change significantly as relaxation need longer time. The RMSD values of DNA in protein bound state sharply increase to their corresponding average values. The root-mean-square deviations (RMSD) of the structures as a function of time with respect to energy minimized structures of the double helices aims to investigate the overall stability of the DNA conformations in protein bound and unbound states during the MD simulations. **Figure S2** does not indicate major structural changes in any system, except the red one (protein unbound operator O2 DNA). That also shows only variations with respect to the average value and no systematic increase as seen in **Figure S1**. Hence, we have analysed the snapshots from the simulation trajectories from 50 ns to 300 ns time steps as RMSD of the structures increase up to 50ns for further studies.

**Figure 2.**
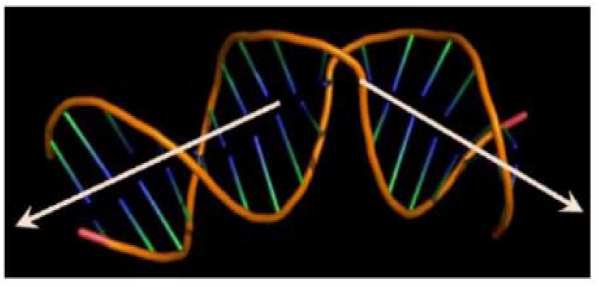
A representative DNA stretch taken from (PDB ID: 1EFA) containing kink. The helix axis bending angle representation is shown by black axis.

**Figure 3:**
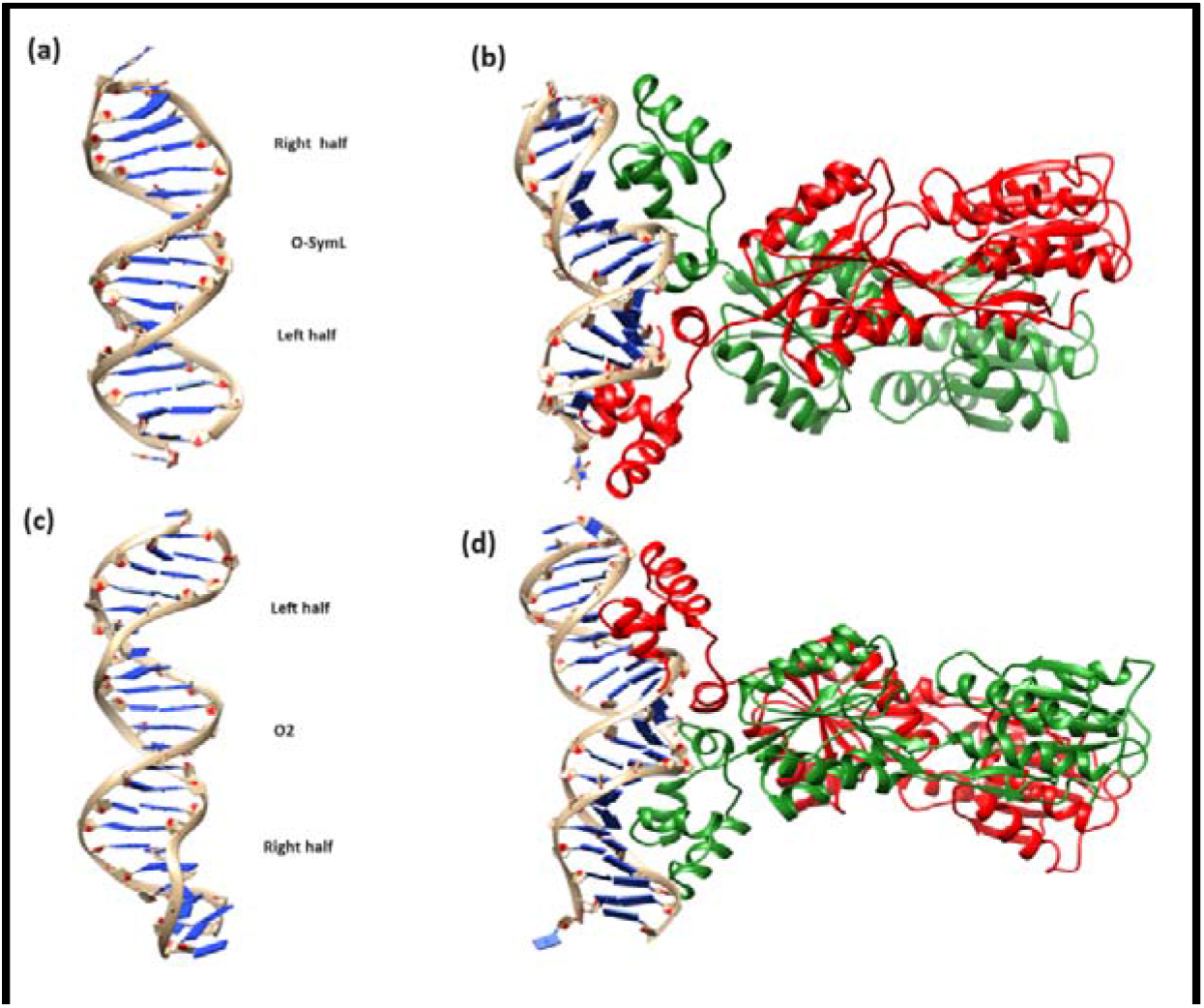
Final snapshot derived from molecular dynamics trajectory for (a) lac repressor unbound operator O-SymL (b) lac repressor bound operator O-SymL (c) lac repressor unbound operator O2 (d) lac repressor bound operator O2

### 3.2 Salient Features of Hydrogen bonding interactions between protein and DNA during MD simulation

The analysis of the hydrogen bonds in Lac-O-SymL and Lac-O2 complexes indicates few amino acid residues make hydrogen bonds and van der Waals contacts with DNA bases but all the hydrogen bonds are not static and maximum numbers of H-bonds are formed between phosphate groups of DNA and amino acid side chains **(Table S1 (a) and (b))**. Alterations in H-bonds involving sugar-phosphate are prominent in the right half-site of the both operators. The hydrogen bonds involving Gln18 (A), which is consistent in the consensus half of O-Sym system with 92.5% stability, disappear in the consensus half of the O2 system. Some hydrogen bonds are, however, exclusively seen in the O2 system, such as between Thr34 (A), Ser31 (A) and sugar-phosphate group of 4 dT of 1^st^ strand as well as Gln60 (A) and sugar-phosphate group of 15 dC at 2^nd^ strand.Thr-34 B(OG1)…(O1P/O2P) Thy-4E H-Bond interaction appears in chain B but disappears in chain A in the O-Sym system. Expected symmetric dynamics is broken in the Lac repressor-symmetric operator complex.

Strong frequent hydrogen bonds Tyr-17A (OH)…(O6) Gua-7D, Gln-18A (NE2)…(O6) Gua-5D, Tyr-17B (OH)…(O6) Gua-7E, Arg-22B (NH1)…(N7) Gua-5E, Arg-22B (NH2)…(O6) Gua-5E occur between Lac repressor protein side chains and base atoms of operator O-Sym complex during MD simulation with more than 90% occupancy. Tyr-7A (OH)…(N4) Cyt-13E, Tyr-17A (OH)…(O6) Gua-7D, Arg-22B (NH1)…(N7) Gua-5E, Arg-22B (NH2)…(O6) Gua-5E H-Bond attribute highest stability for Lac-O2 system. Some of the H-Bonds exist exclusively in the crystal structure of Lac repressor-operator O-Sym complex but disappears in MD simulation e.g., Gln-18A (NE2)…(N4) Cyt-15E, Arg-22A (NH2)…(O6/N7) Gua-5D, Tyr-47A (OH)…(O1P) Cyt-13E, Asn-50A(N)…(O1P) Cyt-13E, Gln-18B(NE2)…(N4) Cyt-15E, Thr-19B(OG1)…(O2P) Gua-5E, Arg-22B(NH2)…(O4) Thy-4E, Arg-22B(NH2)…(N7) Gua-5E, His-29B (ND1)…(O2P) Thy-4E, Ser-31B (N)…(O1P) Thy-4E, Tyr-47B (OH)…(O1P) Cyt-13D. The absence of the central GC base pair in the symmetric operator and the intrinsic asymmetry of the two half-sites in the natural operator O2 lead to significant differences in the binding mode of Lac repressor to the different operator sequences. In addition to the HTH motif, several residues (Tyr47, Asn50 and Gln54) from neighbouring regions of the repressor, are involved in H-bond interaction with the DNA.

### 3.3 Change in dynamics of the protein side chains due to binding with the operator DNA

**Figure S3** compares RMSF of Cα atoms of two individual subunits of Lac repressor with two different force fields Amber-ff99SB-ILDN and Amber-parmbsc1 to ensure that the reported AMBER results are not simulation artifacts. Significant asymmetric dynamics at the protein–DNA interface are observed in both Lac-SymL complex and Lac-O2 complex as well as in the operator unbound state.

In the case of the RMSF plot of DNA SymL bound with Lac repressor, we observe that there is least fluctuation in DNA bases which make hydrogen bonds with amino acids. The differences in dynamics are enhanced in some residues (29His, 31 Ser, 32Ala, 46Asn, 58Gly, 59Lys, 60Gln, 61Leu, 101Arg, 102Ser, 103Gly, 142Asn, 236Gly, 237 Ile, 310Gly, 311Gln, 312Ala, 313Val and 314Lys). Interestingly, the most enhancement of dynamicity upon operator SymL and O2 binding occurs in the hinge and turn and coil conformation. In the operator unbound form the loop showed enhanced mobility, which becomes more rigid upon binding to specific DNA. Both monomers of Lac repressor protein insert hydrophobic side chain Leu56 into the minor groove of DNA as well as unstack two contiguous central base pairs CpG producing a kink at this site. This van der Waals interaction is extremely stable and is well maintained throughout all of the simulation runs. Reduced RMSF value of 56 Leu of Lac repressor in operator bound state (SymL bound A: 1.029 Å, B: 1.0Å and operator O2 bound A: 0.943 Å, B: 0.848 Å) compared to the free state (operator unbound A: 1.57 Å, B: 1.62 Å) also account the phenomena. Noticeably, many N and C-terminal core domain residues of Lac-O2 complex show enhanced fluctuations in the subunit chain A (for example, around residues 75-230) **(Figure 4)**. No significant alteration of dynamics is observed in the case of the Lac repressor-SymL complex. Asymmetric dynamics for protein dimer in operator unbound state is pronounced around residues 285-310 far distant from protein-DNA interface. We reach a conclusion that operator O2 bound Lac repressor shows a distinctly different dynamic character from operator SymL bound complex in chain A. Difference in overall protein dynamics upon interaction with DNAs of different sequences is perhaps the key feature in proper interaction of two C-terminal domains during tetramerization of the Lac repressor dimers for its biological function.

**Figure 4:**
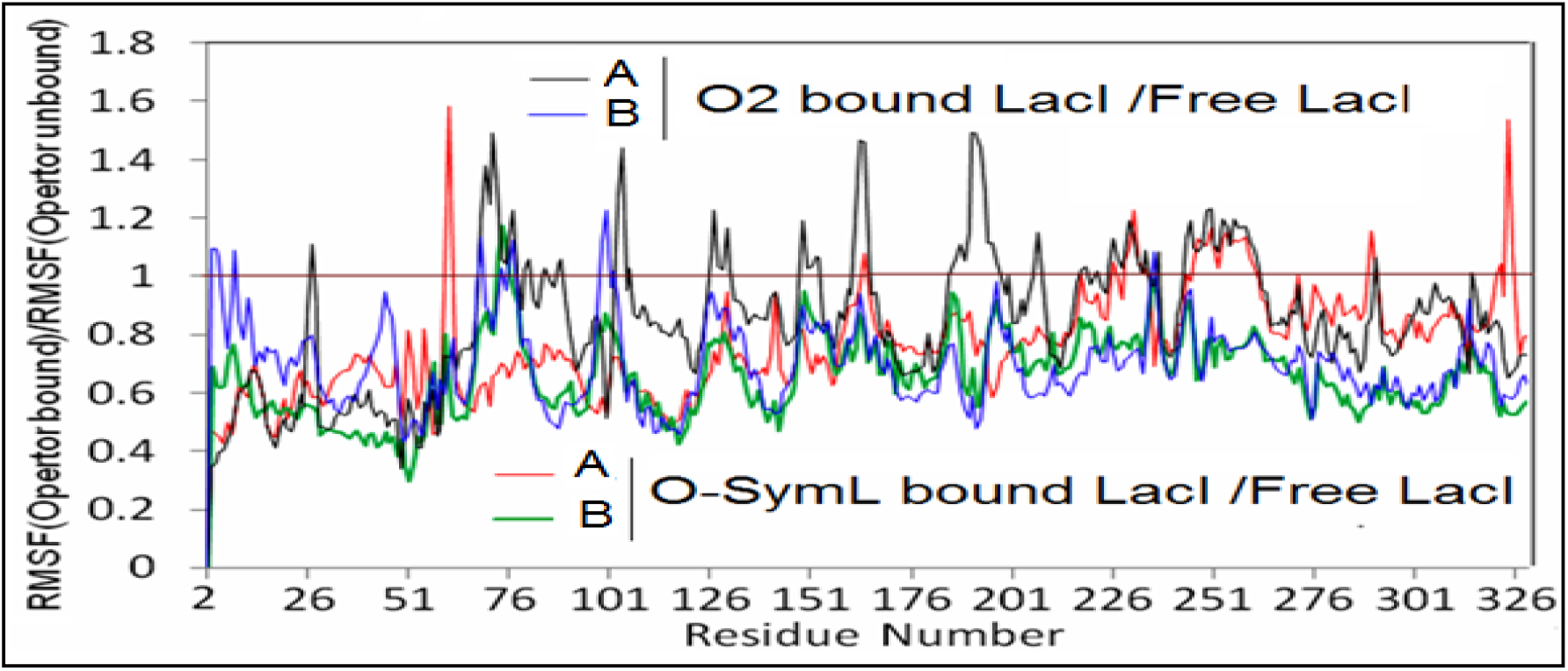
Ratio of RMSF of C_α_ atoms of individual subunits of Lac repressor in absence and presence of operator DNA for both the chains of Lac repressor dimer in operator bound and unbound states with Amber-ff99SB-ILDN force field.

Palindromic, symmetric and synthetic SymL Lac operator with highest binding affinity shed light on symmetric dynamics of both left and right consensus half **(*Figure S4 (a)*)** in protein bound state. We observe asymmetric dynamics (Protein bound strand 2) in the non-consensus half of operator O2 in the protein bound state **(*Figure S4 (b)*)** which is not inconsistent with mutational analysis **[18, 19]**. The nucleotide flexibility reflected by the RMSF values depends on the relative position in the double-stranded DNA chains. Large flexibility at terminal residues of DNA is akin to the end fraying observed in duplex DNA. DNA dynamics decreases in the presence of the protein. The protein clearly reduces the mobility of the phosphate groups of DNA within the binding site and the effect is particularly strong for the DNA backbone involved in hydrogen bonds as well as van der Waals interaction.

### 3.4 Binding thermodynamics from a histogram of side chain dihedral χ1

The equilibrium histogram of structural parameters, such as torsion angles, provides the probability of finding the system in a given conformation, where they can be associated with the corresponding effective free energies with the Boltzmann factors. The Gibbs theory **[20]** provides entropy cost due to conformational changes of the protein. Side chain dihedral χ1 is used as a microscopic conformational variable. Previous studies have estimated thermodynamic parameters from histograms of conformational variables **[21, 22]**. We generated histograms corresponding to χ1 for both the protein bound and unbound states, from the trajectories. In **(Figure 5 (a) and (b))** we observe in addition to single peak histograms, multimodal distributions which clearly indicate different rotameric states corresponding to different fluctuations of some representative amino acids.

**Figure 5:**
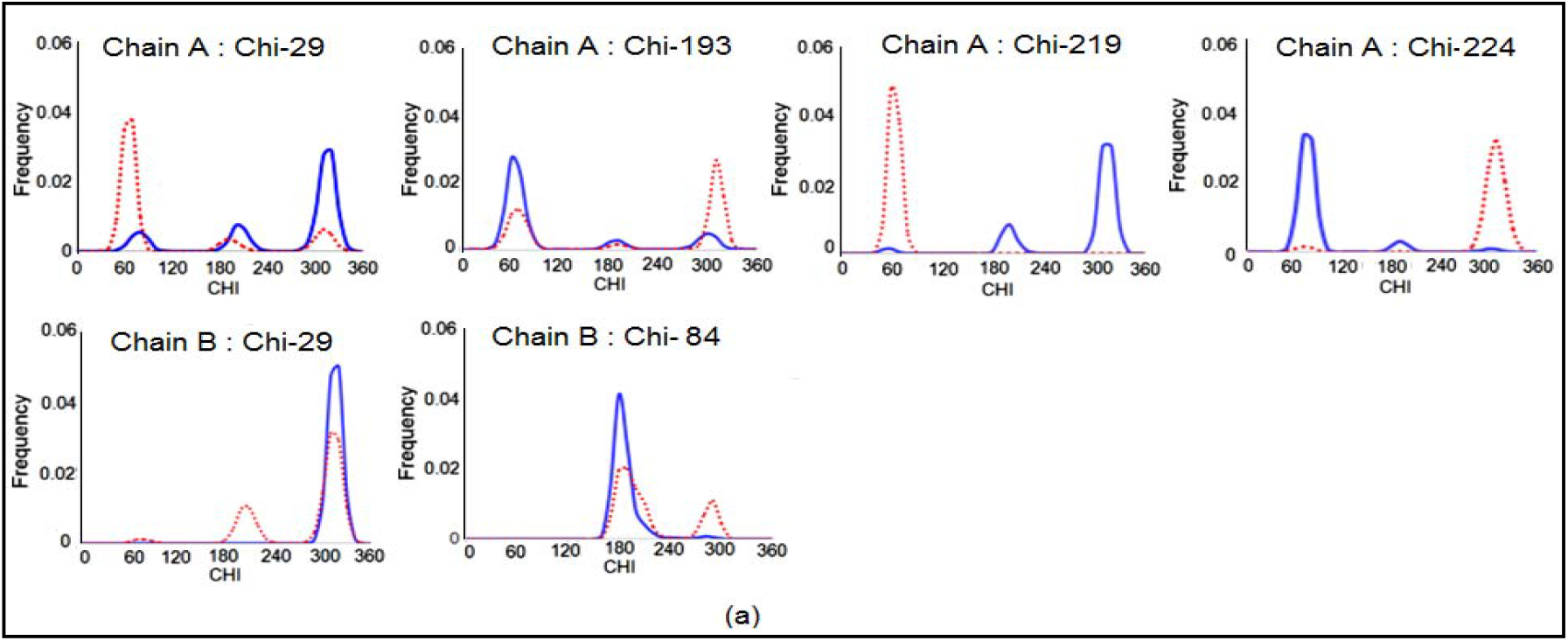
Multimodal histogram of side chain torsion angle, _χ_1 for few protein residues of individual subunits of Lac Repressor in absence (Red dotted line) and presence (Blue line) of operator DNA O-Sym (a) chain A: His29, Ser 193, Asp 219, Ser224; chain B: His29, and Lys 84

**Figure 5:**
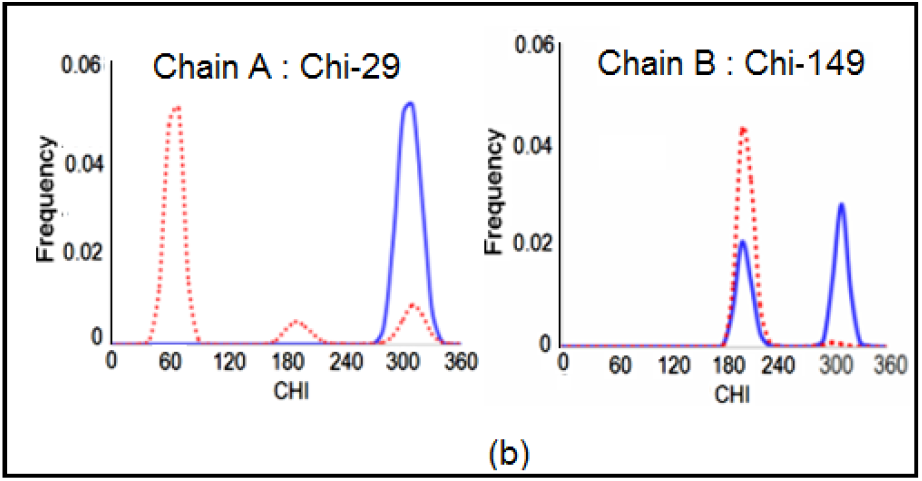
(b) Operator DNA O2 chain A (His29) and chain B (Asp 149) indicating different rotameric states.

Most of the residues of both the chains in the long hinge or loop region (His 29, Asp 149 and Asp 219) have larger fluctuation in terms of conformational entropy. Most of the N-terminal residues (DNA binding domain) of both dimers in Lac repressor-O-SymL show a reduction in entropy due to stable sequence specific H-bond with DNA. But few residues of the DNA binding loop region in contrast show enhancement of conformational entropy in the operator bound state. Conformational entropy of the protein residues in the HTH region is higher in the case of the protein subunit which binds to non-consensus half than the protein subunit that binds to the consensus half for pseudo-palindromic operator bound state. Asymmetric motional freedom is pronounced in the Lac-O2 complex. Many residues in the N and C-terminal sub domains around the protein–protein interaction site show enhanced entropy values **(Figure 6)**. In the operator bound state, augmented conformational entropy is observed in many residues of the N and C-terminal core domain of both the protein chains at or around the protein–protein interaction site. Residue His-29 and Ser-224 show most enhancement of fluctuation in the DNA bound complex. We observe a pronounced enhancement of fluctuation in residue Tyr-7, Ser-16, Gln-18, Val-38, Gln55, Leu63, Val-111 and Ile-213 for the Operator O2 bound complex.

**Figure 6:**
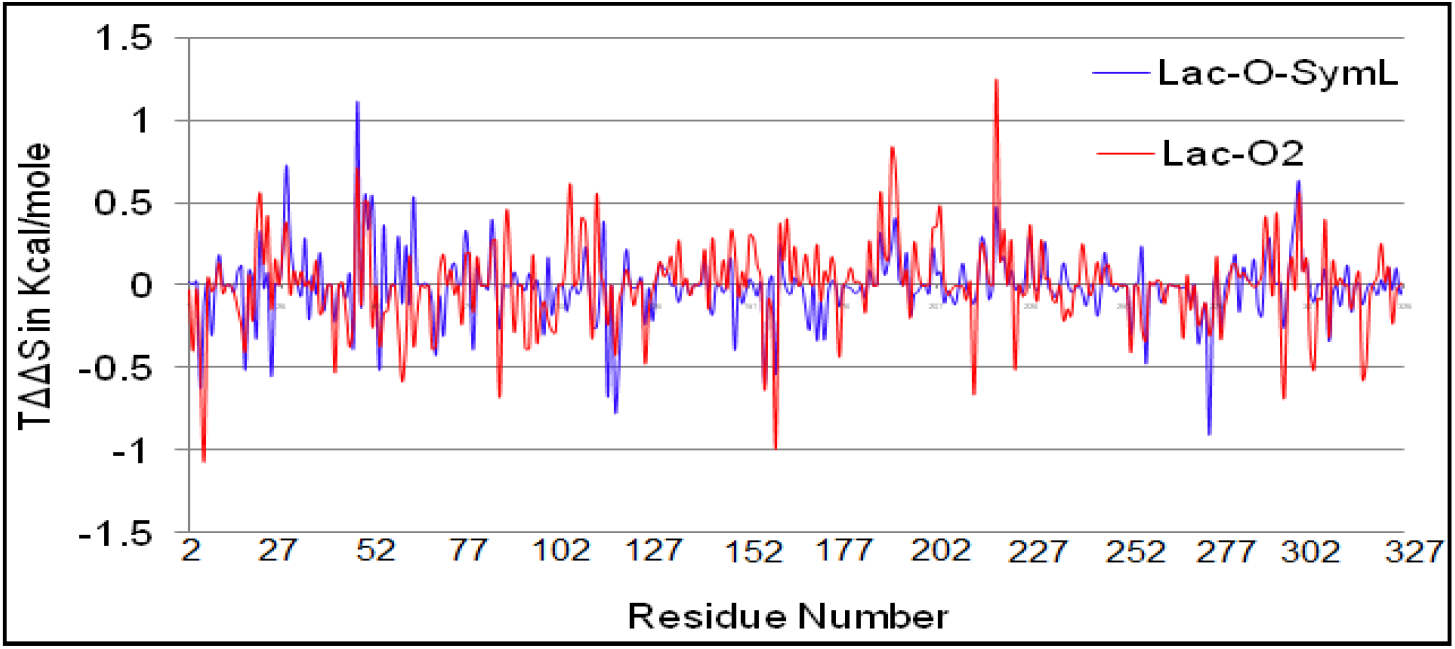
Residue wise conformational entropy change.

Being a homodimer with an identical primary sequence, asymmetric motional freedom of different subunits is unexpected in Lac repressor. We find disorder in loop conformation between helices II and III although there exists a rigid backbone for residues in the three alpha-helices and for the turn of the helix-turn-helix motif. A significant decrease in flexibility is observed for DNA-contacting side chains upon complexation. Most of the N-terminal residues of both dimers in Lac repressor-SymL complex show a reduction in entropy due to additional interaction with the DNA. Residue-wise conformational entropy changes suggest ordering of the residues which form quite a stable sequence-specific H-bond with DNA. Further study of their domain motions and essential dynamics of the protein subunits can shed light on this aspect. Few residues of the DNA binding region also show enhancement of conformational entropy in the operator bound state. Conformational entropy of the protein residues in the HTH region is higher in case of the protein subunit which binds to the non-consensus half than the protein subunit binding to the consensus half for pseudo-palindromic operator bound state. Thus, an important conclusion may be drawn that some residues at or near the protein–protein and protein-DNA interface attain enhanced mobility upon DNA binding.

### 3.5 Conformational transitions and Dynamic Domain Motions study through PCA

Correlated motions in bio-molecules play a pivotal role in mechanical/thermodynamic energy transport and allosteric signal transduction. Since many functional processes involve large and slow conformational changes (as opposed to small-amplitude fast thermal vibrations), the conformational transitions of the complexes are investigated here by projecting their trajectories onto a two-dimensional subspace spanned by the first eigenvectors (PC1) as the first principal component captures the maximum variance in the data set.

The Porcupine plots describe the sub domain motion of Lac-O-SymL, Lac-O2 protein-DNA complex and operator unbound free protein. The 1^st^ principal component for free Lac repressor protein confirms the observation made from the RMSF plot that the most prominent motions are observed at the helix-turn-helix motif of the DNA binding domain of both monomer A and B **(Figure 7)**.

**Figure 7:**
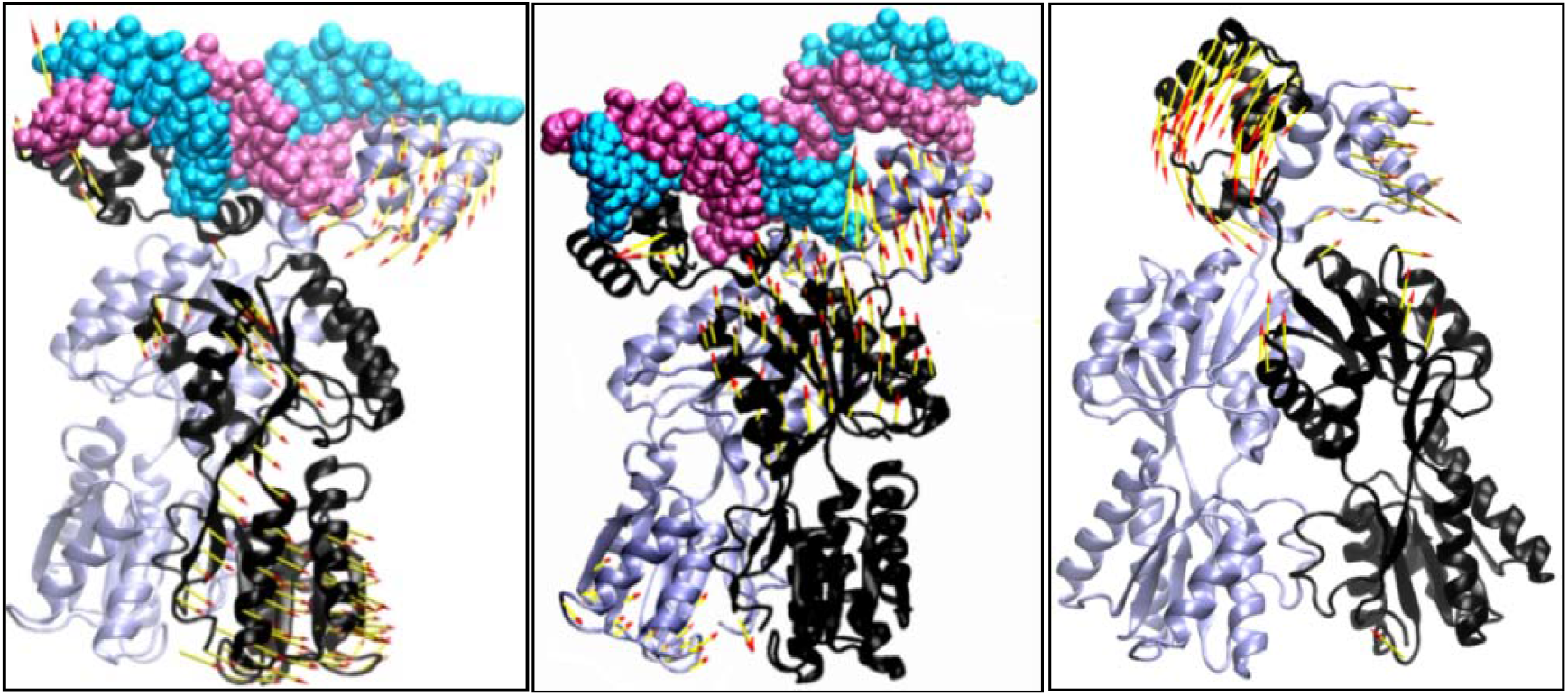
Porcupine plots of the PC1 for the (a) operator SymL bound Lac repressor protein, (b) operator O2 bound Lac repressor protein and (c) free lac protein. Chain C and D: DNA (Chain C: cyan, Chain D: mauve); Chain A and B: Protein (Chain A: iceblue, Chain B: black) The arrows indicate the direction and magnitude of the motion.

The principal component for the Lac-O-SymL protein-DNA complex indicates that the regulatory domain of monomer B undergoes overall greater motion than that of monomer A. Most prominent motions are observed at the loops connecting the α-helices and DNA binding domain. Porcupine plots for the Lac repressor-O-SymL protein-DNA complex simulation focus on the characteristic movement of each monomer. Most of the motion of protein Lac-O-SymL and Lac-O2 protein-DNA complex is sustained in the loop and turn conformation. Protein subunit A binds the left site and B binds right site of the operator. Domain motions and essential dynamics of the protein chains shed light on the asymmetric dynamics of the two monomers and enhancement of fluctuation in the N- and C-terminal core domains for both the monomers due to binding with the operator O2. The protein subunits tend to move away from each other. In operator unbound state of protein, major dynamics is restricted in DNA binding domain. We find a little dynamics far away from DNA binding domain in free Lac repressor protein (in the absence of operators).

How the allostery relates to the regulation of gene expression or how such effects are transmitted, are not known in general. Essential dynamics which enables large scale correlated motions of atoms in the protein-DNA complex can be identified and extracted from MD trajectories, eliminate recurrent modes of structural changes from sets of structures, uncovering the dominant modes within the movement. Intrinsic asymmetry of the two half-sites in the natural operator O2 and absence of the central GC base pair in the symmetric synthetic operator SymL suggest the possibility of significant differences in principal mode of motion of symmetric dimeric protein Lac repressor after binding to the operator sequences.

In summary, the porcupine plot points out how DNA sequence allosterically modulates the dynamic character of the domain of the protein far away from its DNA-protein interaction site. How the allostery relates to the regulation of gene expression or how such effects are transmitted, is still a big puzzle.

### 3.6 Variation of secondary structure of protein subunits with simulation time

To understand the effect of target DNA binding on the protein dynamics, the Operator DNA unbound structure i.e. the dimeric protein alone (system 3) was also studied. Residues 50-58 are disordered for both subunits in operator unbound state. They lose their helicity. Turn and coil secondary structure predominates during the course of simulation. Residues near 235 of the free protein adopt alpha-helical conformation throughout the simulation time but undergo bent conformation in the DNA bound forms. For both subunits upon binding to the specific sequence of O-SymL DNA, residues 50-58 will fold into an α helix which is called the hinge helix. Residues 50-58 are disordered for both subunits upon binding to operator O2. These residues lose their helicity and get converted to unstable loop regions. φ and ψ angles get altered in this region and conformation is quite different in free and complexed proteins **(Figure S5 (a) and (b))**.

### 3.7 Alteration of structural properties of DNA double helix in Multipartite Operator Recognition by Lac repressor

#### 3.7.1 Essential Dynamics Study of DNA Double Helix

It is a difficult task to observe DNA breathing (spontaneous local conformational fluctuation within double stranded DNA) with available experimental methods using ensemble-averaged properties, like NMR, Fluorescence Correlation Spectroscopy, etc., due to the low frequency and short duration of base pair opening **[23]**. So we explore DNA breathing dynamics by examining inter-base pair parameters, groove width, and bending angle using PCA to investigate whether the conformational changes associated with partial intercalation of two Leu amino acid ligands into DNA are restricted to just the two bases above and below the intercalation site or not. The most prominent anti-correlated rotary motion is observed in the minor groove. The left consensus half of the model moves in the anti-clockwise direction and the right consensus half moves in the clockwise direction in the case of the free O-Sym operator.

#### 3.7.2 Bending and flexibility of DNA

The bending angle of protein unbound operator O-Sym and O2 are much lower (average value ∼15^0^) **(Table 3 and Figure S6)**. Significant overlap between the bending histograms of repressor-operator complexes (Lac repressor-O-SymL and Lac repressor-O2) indicates bending deformability that leads to deformed conformation.

**Table 3:** Mean and SD of Bending Angle in degree (ff corresponds to Amber-ff99SB-ILDN)

| System | Mean | PCA-first | PCA-last |
| --- | --- | --- | --- |
| Operator O-SymL(Complex) | 44.15(5.8) | 39.8 | 45.2 |
| Operator O-SymL(Free) | 16.32(8.9) | 43.8 | 22.2 |
| Operator O2(Complex) | 37.22(5.4) | 47.0 | 32.3 |
| Operator O2(Free) | 14.5(7.5) | 43.2 | 26.1 |

The protein-bound DNA, shows the distribution of bending angle which matches with normal distribution. Both in protein free and protein bound conditions, the Lac-O2 operator appears to have a lesser bending propensity, which may be one of the reasons for the poorer repressing capability of O2 as compared to O-Sym. The free DNA shows low average bending but large difference between bending in PCA-first and PCA-last. The protein bound DNA, however, shows a large mean bending as well as smaller differences in PCA based bending angles. This probably indicates some conformational selection mode of recognition takes place by the DNA.

PCA analysis of bending angle deciphers operator O2 has more propensity to be deformed compared to operator O-SymL. On the other hand, protein bound O-Sym shows relatively little natural breathing of the bending angle. **Table 3** clarifies that free operators travel the conformational spaces in a similar way but both of which are different from the conformational space traversed by the DNA molecule bound with the protein. Principal components of bending angle with two values 39.8 and 45.2, (the difference is very small) for operator O-Sym complex indicates a certain amount of static bending (perhaps due to highest affinity of SymL operator for Lac repressor) while the comparatively large difference between principal components of bending angle indicates dynamic bending or flexibility for free operator O-Sym and O2 as well as the Operator O2 complex.

#### 3.7.3 PCA of Groove Width in terms of P-P Distance

At the centre of the operator DNA, the minor groove width (P-P distance) of the binding region of DNA in the complex is found to be large and their standard deviations are also significantly less as compared to other free DNA systems. Significant breathing is observed for both major and minor grooves of all protein bound and free operators except the minor groove of protein bound O-Sym. Remarkable variations in minor groove widths at the intercalated site arise from differences in the hydrogen bonding pattern of each base pair and from different stacking interactions for each di-nucleotide step **(Figure S7)**.

#### 3.7.4 Base pair dynamics of DNA during MD simulations

The overall analysis of roll, slide and Zp parameters of protein unbound operators confirms the simulated DNA structures to be in B-form. The roll value of kinked central CG/CG in both Lac-O-SymL operator and Lac-O2 operator is highest and twist value for this corresponding step is lowest due to partial intercalation of 56 Leu of the hinge region of Lac repressor in these two base pair steps. The rise values, i.e., the separation between the two base pairs at this step, increase from 3.4 Å to 3.8 Å; hence we can conclude that total intercalation of amino acid side chains does not occur. DNA untwisting results in an increase of the minor groove size and a narrower major groove compared to standard B-DNA. Significant reduction of overlap values also accounts for unstacking at the kinked central CpG step. δ torsion angles in these central bases are in the 90° range at one of the strands, corresponding to mixed puckering pattern of C3’-endo sugar pucker at one strand and C2’-endo sugar pucker at another strand. Rigidity of residues of protein in the binding side may also help to hold the DNA in a bent conformation.

## 4. Conclusion

Despite being a symmetric homodimer and having secondary structure similarity, two protein subunits bound to O-Sym and O2 as well as protein unbound are dynamically non-equivalent. Asymmetry of protein-DNA structure and dynamics in presence of sequence identity is key feature for the O-Sym system. The difference of hydrogen bonding network and deformation of DNA base pair step produce the asymmetric dynamics of the two subunits of Lac repressor protein. Variation of minor groove widths can direct overall shape complementarily between the DNA and protein surfaces that means DNA is significantly deformed to accommodate the protein fold. Bending flexibility indirectly modulates the Lac binding activity that controls transcription efficiency. Modulation of protein–protein interactions by the different DNA sequences stabilize the gene regulatory networks differently and perhaps attribute in recognising two C-terminal domains during tetramerization of the Lac repressor dimers for its biological function. Asymmetric dynamics of two Lac repressor protein subunits bound to O-Sym and O2 points out the role of DNA sequence specific interaction which is very important in the context of transcription regulation. Lac operator is unique as it uses both Lock & Key mechanism for DNA recognition, where protein induced structural alteration of the receptor DNA is not required, and also Induced Fit/Conformational Selection model of recognition.

## Acknowledgments

Soumi Das is thankful to the CSIR-NET fellowship for financial support.

