## Supplementary Information for "In-silico studies on structural and thermodynamics basis of interaction of lac repressor protein binding to different DNA operators"

**^1^Department of Biophysics, Bose Institute, Kolkata 700054;**

**^2^Computational Science Division, Saha Institute of Nuclear Physics, Kolkata 700064**

**Correspondence: Soumi Das**

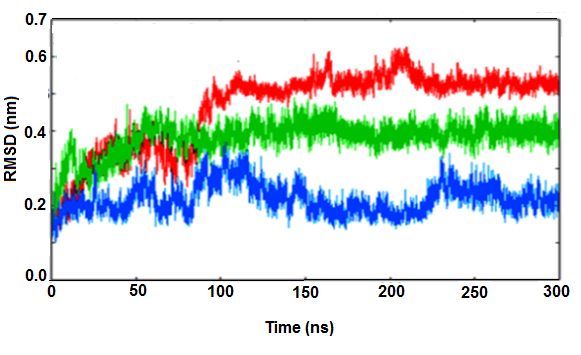

***Figure S1: RMSD of the trajectories Lac repressor protein in operator SymL bound complex (system 1 in blue), Lac repressor protein in operator O2 (system 2 in green) bound complex and operator unbound free Lac repressor protein (system 3 in red) calculated with respect to the energy minimized structure considering C-alpha atom of Protein***

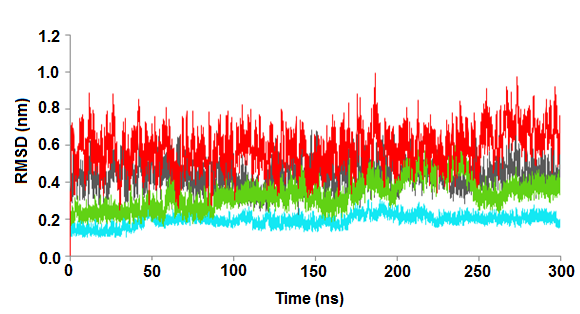

***Figure: S2: RMSD of the trajectories using Amber-ff99SB-ILDN ff: DNA Operator SymL DNA (Protein bound) in Cyan, Operator O2 DNA (Protein bound) in Green, Operator SymL DNA (Protein unbound) in Black, Operator O2 DNA (Protein unbound) in Red***

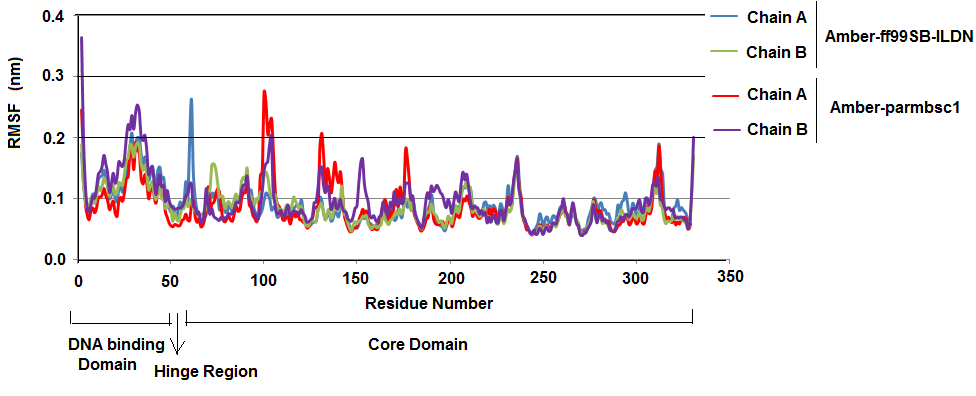

***Figure S3: Comparison of RMSF of Cα atoms of individual subunits of Lac chain A and chain B with two different force field (Amber-ff99SB-ILDN and Amber-parmbsc1)***

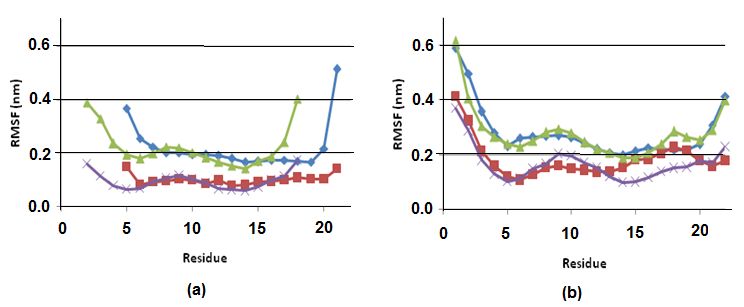

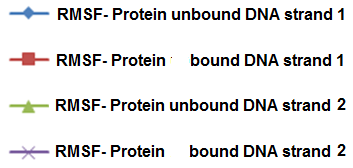

***Figure S4: RMSF of phosphorus atoms within the isolated DNA and DNA-Protein complex of (a) operator O-Sym (b) operator O2***

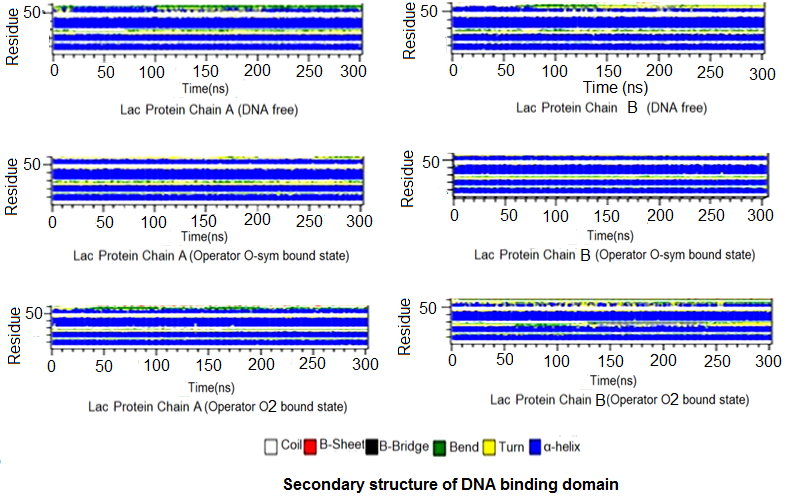

***Figure S5: (a) Variation of secondary structure of protein subunits DNA Binding domain with simulation time***

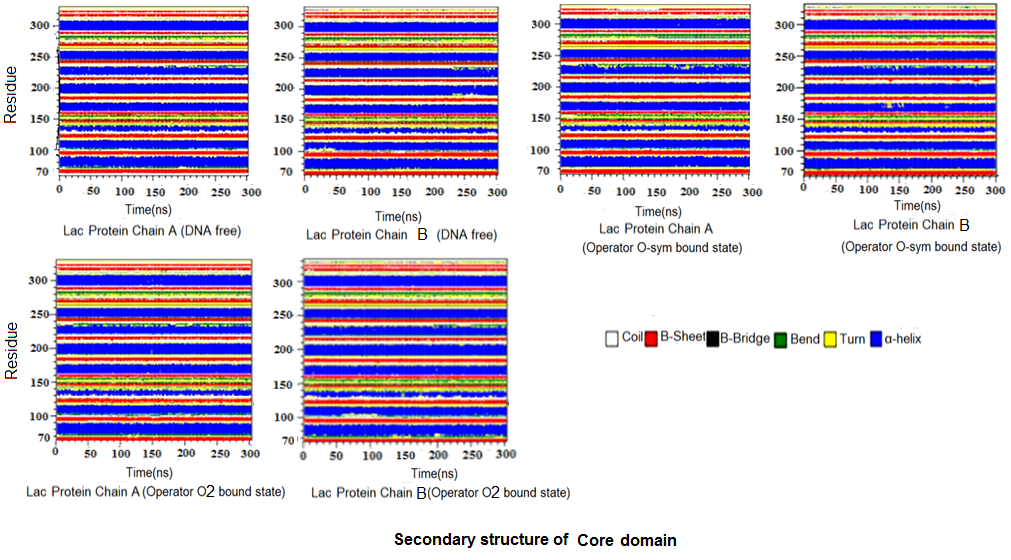

***Figure S5: (b) Variation of secondary structure of protein subunits Core domain with simulation time***

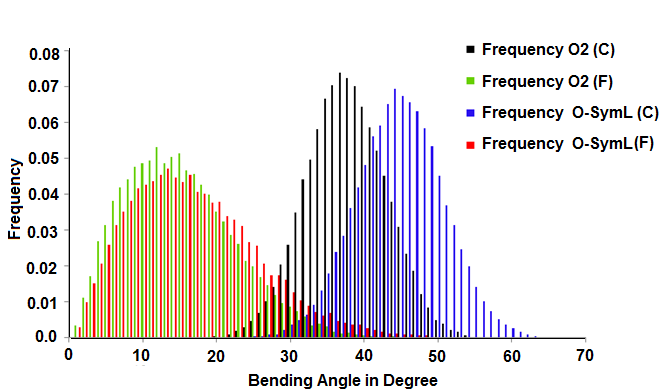

***Figure S6: Distribution of bending angle for free DNA of SymL (red) and in the SymL (complex) (blue) as well as free DNA of O2 sequence (green) and in the O2 (complex) (black) indicates bending deformability of DNA.***

P-P Minor groove width
in A

|  | PC1O-SymL(C)1st | PC1O-SymL(C)Last | PC1O-SymL(F)1st | PC1O-SymL(F)Last |
| --- | --- | --- | --- | --- |
| 8DA-17DC | 14.6 | 13.6 | 16.71 | 16.22 |
| 9DG-16DA | 11.57 | 12.47 | 11.76 | 12.31 |
| 10DC-15DC | 13.4 | 13.06 | 11.33 | 12.89 |
| 12DG-14DT | 16.17 | 16.88 | 12.54 | 12.63 |
| 13DC-13DC | 19.75 | 18.42 | 14.87 | 14.61 |
| 14DT-12DG | 15.95 | 15.45 | 13.78 | 11.25 |
| 15DC-10DC | 12.54 | 13.6 | 13.98 | 11.32 |
| 16DA-9DG | 12.74 | 13.99 | 15.43 | 12.12 |
| 17DC-8DA | 13.59 | 14.06 | 16.1 | 15.32 |
| 18DA-7DG | 12.26 | 12.62 | 14.14 | 14.57 |
| 19DA-6DT | 11.46 | 11.23 | 13.41 | 14.03 |
| 20DT-5DG | 11.86 | 12.16 | 11.8 | 13.22 |

***Figure S7: (a)***

P-P Minor groove width
in A

|  | PC1O2(C)1st | PC1O2(C)Last | PC1O2(F)1st | PC1O2(F)Last |
| --- | --- | --- | --- | --- |
| 5DG-21DT | 10.73 | 12.75 | 13.04 | 10.74 |
| 6DT-20DT | 10.73 | 9.78 | 12.73 | 12.26 |
| 7DG-19DT | 9.9 | 12.92 | 14.38 | 12.62 |
| 8DA-18DA | 12.36 | 11.37 | 18.59 | 14.65 |
| 9DG-17DC | 12.6 | 13.11 | 19.42 | 13.55 |
| 10DC-16DA | 13.13 | 12.6 | 13.49 | 10.6 |
| 11DG-15DC | 15.07 | 13.59 | 12.99 | 10.33 |
| 12DA-14DT | 20.25 | 19.47 | 15.27 | 12.28 |
| 13DG-13DC | 19.55 | 20.42 | 12.06 | 12.39 |
| 14DT-12DG | 15.48 | 14.97 | 10.61 | 12.86 |
| 15DA-11DC | 15.78 | 9.16 | 9.32 | 12.65 |
| 16DA-10DT | 11.11 | 16 | 11.12 | 15.24 |
| 17DC-9DC | 13.13 | 22.49 | 12.46 | 14.73 |
| 18DA-8DA | 9.67 | 16.12 | 13.37 | 14.14 |
| 19DA-7DT | 11.48 | 11.45 | 16.53 | 13.53 |
| 20DC-6DT | 12.26 | 10.27 | 15.98 | 13.18 |
| 21DC-5DG | 11.81 | 11.15 | 12.3 | 11.88 |
| 22DG-4DT | 10.96 | 10.52 | 12.12 | 11.34 |

***Figure S7: (b)***

***Figure S7: Minor groove width as distance between P-P***

***(1^st^ and Last mean two end points of vibration as obtained from porcupine plot)***

***C = Operator in protein bound and F = Operator in protein free***

***Table S1: (a) Hydrogen bonds***

(Donor to Acceptor Distance ≤ 3.5 Ǻ and H-bond angle ≥ 120^0^)

For DNA, chain D is 1^st^ strand and chain E is 2^nd^ stand

***(Table for 1EFA using pyrHBfind and pdb file numbering follows Figure 1)***

***Hydrogen bonds between Lac repressor and operator O-Sym from MD trajectory (50ns-300ns)***

| **Donor** | **Donor Atoms** | **Acceptor** | **Acceptor Atoms** | **% of Occurrence** | **Hydrogen bond distance**  **in Crystal** |
| --- | --- | --- | --- | --- | --- |
| Lys-2A | NZ | Gua-12E | O2P | 48.61 |  |
| Leu-6A | N | Cyt-13E | O2P | 100 | 2.22 |
| Tyr-7A | OH | Cyt-13E | N4 | 70.0 | 2.46 |
| Ser-16A | N | Gua-5D | O1P | 53.56 | 2.92 |
| Ser-16A | OG | Gua-5D | O2P | 99.91 | 1.99 |
| Ser-16A | N | Gua-5D | O2P | 99.84 |  |
| Ser-16A | OG | Gua-5D | O5’ | 18.61 |  |
| Tyr-17A | OH | Gua-7D | O6 | 99.36 | 1.94 |
|  | OH | Gua-7D | N7 | 74.64 | 2.85 |
| Gln-18A | NE2 | Cyt-15E | N4 |  | 2.42 |
| Gln-18A | NE2 | Gua-5D | O6 | 92.51 |  |
| Thr-19A | OG1 | Gua-5D | O2P | 99.99 | 2.79 |
| Ser-21A | OG | Thy-14E | O2P | 99.98 | 1.60 |
|  | OG | Thy-14E | O5’ | 55.88 |  |
| Arg-22A | NH2 | Gua-5D | O6 |  | 2.95 |
|  | NH2 | Gua-5D | N7 |  | 2.00 |
|  | NH2 | Gua-5D | O6 |  | 2.83 |
| Asn-25A | ND2 | Thy-14E | O1P | 77.60 | 1.79 |
|  |  | Thy-14E | O2P | 54.60 |  |
|  |  | Thy-14E | O5’ | 66.13 |  |
| Tyr-47A | OH | Cyt-13E | O1P |  | 1.60 |
|  | OH | Cyt-13E | O2P | 99.92 |  |
|  | OH | Cyt-13E | O5’ | 30.94 |  |
| Asn-50A | N | Cyt-13E | O1P |  | 2.52 |
|  | N | Cyt-13E | O5’ | 96.25 |  |
| Gln-54A | NE2 | Thy-14E | O1P | 99.28 | 1.83 |
|  | NE2 | Cyt-13E | O3’ | 23.36 |  |
| Lys-59A | NZ | Cyt-15E | O1P | 56.55 |  |
|  | NZ | Thy-14E | O1P | 19.67 |  |
| Lys-2B | NZ | Gua-12D | O2P | 44.91 |  |
| Leu-6B | N | Cyt-13D | O2P | 99.94 | 2.19 |
| Tyr-7B | OH | Cyt-13D | N4 | 43.64 |  |
| Ser-16B | N | Gua-5E | O1P | 12.91 | 2.53 |
|  | OG | Gua-5E | O2P | 99.62 | 1.6 |
|  | N | Gua-5E | O2P | 99.96 |  |
|  | OG | Gua-5E | O5’ | 74.24 |  |
| Tyr-17B | OH | Gua-7E | N7 | 72.8 | 2.85 |
|  | OH | Gua-7E | O6 | 99.66 | 2.21 |
| Gln-18B | NE2 | Cyt-15D | N4 |  | 2.33 |
| Gln-18B | NE2 | Thy-6E | O4 | 28.86 |  |
| Thr-19B | OG1 | Gua-5E | O2P |  | 2.58 |
|  | OG1 | Gua-5E | O2P | 99.83 |  |
| Ser-21B | OG | Thy-14D | O2P | 15.5 | 1.60 |
| Arg-22B | NH2 | Thy-4E | O4 |  | 2.50 |
|  | NH2 | Gua-5E | O6 | 99.93 | 2.49 |
|  | NH2 | Gua-5E | N7 |  | 2.19 |
|  | NH1 | Gua-5E | N7 | 98.34 |  |
|  | NH1 | Gua-5E | O6 | 32.86 |  |
| Asn-25B | ND2 | Thy-14D | O1P | 10.76 |  |
|  | ND2 | Thy-14D | O2P | 24.5 |  |
| His-29B | ND1 | Thy-4E | O2P |  | 2.25 |
| Ser-31B | N | Thy-4E | O1P |  | 2.31 |
|  | N | Thy-4E | O2P | 99.76 | 2.78 |
|  | OG | Thy-4E | O1P | 50.86 |  |
|  | OG | Thy-4E | O2P | 17.49 |  |
| Thr-34B | OG1 | Thy-4E | O1P | 95.77 | 1.60 |
|  | OG1 | Thy-4E | O2P | 17.80 |  |
| Tyr-47B | OH | Cyt-13D | O1P |  | 1.81 |
|  | OH | Cyt-13D | O2P | 33.42 |  |
| Asn-50B | ND2 | Cyt-13D | O1P | 68.00 |  |
|  | N | Cyt-13D | O5’ | 30.59 |  |
|  | ND2 | Gua-12D | O5’ | 15.1 |  |
| Gln-54B | NE2 | Thy-14D | O1P | 95.9 | 2.01 |
| Lys-59B | NZ | Thy-14D | O1P | 37.43 |  |

***Table S1: (b) Hydrogen bonds between Lac repressor and operator O2 from MD trajectory (50ns-300ns)***

| **Donor** | **Donor Atoms** | **Acceptor** | **Acceptor Atoms** | **% of Occurrence** |
| --- | --- | --- | --- | --- |
| Lys-2A | NZ | Gua-12E | O2P | 32.10 |
| Leu-6A | N | Cyt-13E | O2P | 99.93 |
| Tyr-7A | OH | Cyt-13E | N4 | 97.48 |
| Ser-16A | N | Gua-5D | O2P | 99.96 |
|  | OG | Gua-5D | O2P | 99.84 |
| Tyr-17A | OH | Gua-7D | O6 | 99.31 |
|  | OH | Gua-7D | N7 | 78.58 |
| Thr-19A | OG1 | Gua-5D | O2P | 99.66 |
| Ser-21A | OG | Thy-14E | O2P | 84.66 |
| Arg-22A | NH1 | Gua-5D | N7 | 78.32 |
|  | NH2 | Gua-5D | O6 | 84.56 |
| Asn-25A | ND2 | Thy-14E | O1P | 78.88 |
|  | ND2 | Thy-14E | O2P | 76.68 |
| His-29A | ND1 | Ade-3D | O2P | 26.23 |
| Ser-31A | N | Thy-4D | O2P | 99.82 |
| Thr-34A | OG1 | Thy-4D | O1P | 98.36 |
| Tyr-47A | OH | Cyt-13E | O2P | 73.5 |
| Asn-50A | N | Cyt-13E | O1P | 91.33 |
|  | N | Cyt-13E | O5’ | 71.06 |
| Gln-54A | NE2 | Thy-14E | O1P | 92.84 |
| Lys-59A | NZ | Thy-14E | O1P | 97.34 |
| Gln-60A | N | Cyt-15E | O1P | 95.32 |
|  | NE2 | Cyt-15E | O1P | 15.74 |
| Arg-118A | NH1 | Gua-11D | O1P | 60.72 |
|  | NH2 | Gua-11D | O1P | 51.07 |
| Leu-6B | N | Ade-12D | O2P | 99.07 |
| Ser-16B | N | Gua-5E | O2P | 99.98 |
|  | N | Gua-5E | O1P | 12.5 |
|  | OG | Gua-5E | O2P | 91.29 |
| Tyr-17B | OH | Gua-13D | N7 | 56.12 |
|  | OH | Ade-12D | N9 | 12.82 |
| Gln-18B | NE2 | Thy-6E | O4 | 88.86 |
|  | NE2 | Ade-15D | N7 | 19.32 |
| Thr-19B | OG1 | Gua-5E | O2P | 100 |
| Arg-22B | NH1 | Gua-5E | N7 | 99.47 |
|  | NH2 | Gua-5E | O6 | 99.85 |
| Asn-25B | ND2 | Gua-13D | O2P | 58.26 |
| Ser-31B | N | Thy-4E | O2P | 99.84 |
|  | OG | Thy-4E | O1P | 66.89 |
| Thr-34B | OG1 | Thy-4E | O1P | 88.90 |
| Tyr-47B | OH | Ade-12D | O1P | 99.94 |
| Asn-50B | N | Ade-12D | O1P | 96.10 |
| Gln-54B | NE2 | Gua-13D | O1P | 88.56 |
| Lys-59B | NZ | Thy-14D | O1P | 34.74 |
|  | NZ | Gua-13D | O3’ | 18.63 |
